# Increased endothelium activation and leakage do not promote diastolic dysfunction in mice fed with a high fat diet and treated with L-NAME

**DOI:** 10.1101/2023.02.08.527684

**Authors:** Lauriane Cornuault, Pierre Mora, Paul Rouault, Ninon Foussard, Candice Chapouly, Pilippe Alzieu, Alain-Pierre Gadeau, Thierry Couffinhal, Marie-Ange Renault

**Affiliations:** Univ. Bordeaux, Inserm, Biology of Cardiovascular Diseases, U1034, CHU de Bordeaux, F-33604 Pessac, France

**Keywords:** Heart failure, diastolic dysfunction, microvessels, cardiac remodeling, sexual dimorphism

## Abstract

Coronary microvascular disease has been proposed to be responsible for heart failure with preserved ejection fraction (HFpEF) about 10 years ago. However, to date the role and phenotype of the coronary microvasculature has still been poorly considered and investigated in animal models of HFpEF.

**Objective:** To determine whether endothelial dysfunction participates in the development of diastolic dysfunction in mice fed with a high fat diet (HDF) and treated with L-NAME.

**Approach and Results:** At first, we thoroughly phenotyped the coronary microvasculature in this model in male, female and ovariectomized (OVX) female considering the sexual dimorphism associated with this disease. We found that both OVX and non OVX females but not males display increased endothelial activation, leakage, and arteriole constriction upon the HFD + L-NAME regimen while both male and OVX females but not non OVX females develop diastolic dysfunction. With the aim to investigate the role of endothelial dysfunction in the pathophysiology of diastolic dysfunction in OVX female mice, we used Cdon deficient mice. Indeed, we previously demonstrated that endothelium integrity, upon inflammatory conditions, is preserved in these mice. Both OVX Cdh5-Cre/ERT2-Cdon^Flox/Flox^ (Cdon^ECKO^) and Cdon^Flox/Flox^ (Ctrl) female mice were fed with the HFD + L-NAME regimen to induced diastolic dysfunction. As expected, Cdon^ECKO^ mice displayed improved endothelium integrity i.e. decreased endothelium permeability, decreased ICAM-1 expression and decreased infiltration of CD45+ leukocytes in comparison to control mice. However, Cdon^ECKO^ mice displayed cardiac hypertrophy, cardiac fibrosis and increased end diastolic pressure just like control mice. Moreover, we found that cardiac inflammation does not participate in the pathophysiology of HFpEF either by treating OVX female mice with colchicine.

**Conclusion:** Altogether, the data presented in this paper demonstrate that neither endothelium permeability nor endothelial activation or inflammation do participate in the pathophysiology of diastolic dysfunction in mice exposed to HFD+L-NAME.

## INTRODUCTION

A significant proportion of patients with clinical syndrome of heart failure (HF) happen to have a normal left ventricular ejection fraction (EF) referred to as heart failure with preserved ejection fraction (HFpEF) as opposed to patients with reduced ejection fraction (HFrEF). HFpEF is characterized by increased arterial and myocardial stiffness and decreased left ventricular (LV) relaxation which causes increased LV end-diastolic pressure with impaired LV filling. This diastolic dysfunction is associated with abnormal ventricular-arterial coupling, pulmonary hypertension, chronotropic incompetence and cardiac reserve dysfunction ^1^. Notably, HFpEF now accounts for more than 50% of all HF patients and has turned into the most common HF phenotype ^2^. However, in contrast to HFrEF, for which significant advances in understanding etiologies and mechanisms have been made, HFpEF pathophysiology is still poorly understood ^3^. HFpEF patients are generally older, more often females, with a high prevalence of cardiovascular and non-cardiovascular comorbidities, such as obesity, metabolic syndrome, type 2 diabetes mellitus, hypertension, atrial fibrillation or renal dysfunction ^4^. At molecular and cellular level, HFpEF has been associated with cardiac fibrosis, cardiomyocyte hypertrophy and stiffness, microvascular dysfunction, inflammation, decreased bioavailability of nitric oxide (NO) and oxidative stress ^1^. In 2013, based on the fact that vascular and microvascular dysfunction is highly prevalent in HFpEF patients, WJ Paulus and C Tschöpe proposed the following paradigm: cardiovascular risk factors would lead to a systemic low-grade inflammation triggering endothelial cells (EC) dysfunction which would in turn be responsible for cardiac wall stiffening and diastolic dysfunction ^5^. Over the last decades, the causal role of EC dysfunction has been supported by preclinical studies in which conditional KO rodents have been used to demonstrate that modification of EC properties is sufficient to promote diastolic dysfunction. This is the case of Sirtuin 3 ^6 7^ or Sirtuin 1 ^8^ disruption and β3-Adrenergic Receptor ^9^ or PDZ Domain Containing Ring Finger 3 ^10^ overexpression in ECs.

However, it is still actually not known whether EC dysfunction is responsible for diastolic dysfunction in patients or animal models with HFpEF ^11^. Notably, WJ Paulus’s group proposed that decreased NOS3 activity in ECs would decrease NO available to activate soluble guanylate cyclase (sGC) in cardiomyocyte leading to decreased cGMP production, protein kinase cGMP-dependent 1 (PRKG1) activity and subsequently Titin phosphorylation, ultimately leading to cardiomyocyte stiffening ^5^. However, the negative results of clinical trials aiming at restoring NO/sGC/cGMP signaling in HFpEF patients suggest that this pathway may not have a central role in the pathophysiology of HFpEF ^11^. Importantly, endothelial dysfunction does not only design endothelium-dependent vasodilation but any detrimental modification of the endothelial phenotype. Notably, it also designs impaired endothelial barrier properties, endothelial activation, impaired endothelial paracrine activity and impaired endothelial or quiescence.

Accordingly in the present study, we further explored the phenotype of the cardiac microvasculature in the 2-hit model of HFpEF developed by Schiattarella G et al ^12^ in which diastolic dysfunction is induced by a high fat diet associated with a L-NAME treatment. Importantly, considering the sexual dimorphism possibly associated with HFpEF this study has been done in males, females and ovariectomized females. Additionally, we tested whether endothelial dysfunction, especially abnormal endothelial activation and permeability mice does participate in the development of diastolic dysfunction in ovariectomized female mice using endothelial specific cell adhesion associated, oncogene regulated (*Cdon*) deficient mice.

## METHODS

### Mice

C57BL/6J mice were obtained from Charles River laboratories and bred in our animal facility. Cdon floxed (Cdon^tm1c(EUCOMM)Hmgu^) (Cdon^Flox^)^13^ mice were generated at the Institut Clinique de la Souris through the International Mouse Phenotyping Consortium from a vector generated by the European conditional mice mutagenesis program, EUCOMM. Cdon floxed mice were generated under a C57BL/6N background and backcrossed at least 6 times with C57BL/6J mice before they were used in any experiment.

Tg(Cdh5-cre/ERT2)1Rha, (Cdh5-Cre/ERT2) mice^14^ which were a gift from RH. Adams, were maintained under a C57BL/6J background.

Cdh5-Cre/ERT2 mice were genotyped using the following primers: 5′- TAAAGATATCTCACGTACTGACGGTG-3′ and 5′-TCTCTGACCAGAGTCATCCTTAGC-3′ that amplify 300 bp of the Cre recombinase sequence. Cdon floxed mice were genotyped using the following primers: 5′- CTTCCCAGAGGGTGTGAGAGCAATG-3′ and 5′-GAACCAGTAGCATGCATGATGCTGG-3′ which amplifies a 385 bp fragment of the WT allele or a 494 bp fragment if the allele is floxed.

The Cre recombinase in Cdh5-Cre/ERT2 mice was activated by intraperitoneal injection of 1 mg tamoxifen (Sigma-Aldrich) for 5 consecutive days at 8 weeks of age. To induce Cre, tamoxifen is injected IP 1 mg/mL 5 consecutive days at 8 weeks of age. Successful and specific activation of the Cre recombinase have been verified before ^13^.

Animal experiments were performed in accordance with the guidelines from Directive 2010/63/EU of the European Parliament on the protection of animals used for scientific purposes and approved by the local Animal Care and Use Committee of Bordeaux University. Mice were either sacrificed by cervical dislocation or exsanguination under deep anesthesia (ketamine 100 mg/kg and xylazine 20 mg/kg, IP).

### Ovariectomy

Bilateral ovariectomy was performed at 7-8 weeks of age after anesthesia with 3% isoflurane. To minimize pain caused by the surgery, mice were subcutaneously administered 0.1 mg/kg buprenorphine 30 min before surgery.

### HFD regimen/L-NAME treatment

Mice had unrestricted access to high fat diet containing 60% of the energy from fat (butter) (SAFE^®^ 260 HF) for 12 weeks. Alternatively, mice were fed with regular chow diet (SAFE^®^ A04).

L-NAME (Sigma Aldrich) was supplied in the drinking water at 1 mg/L concentration 4 days/7 for 10 weeks.

### Treadmill Exercise Exhaustion Test

Before treadmill experiments, mice were habituated to handling to reduce anxiety. Then, animals were familiarized with running on the treadmill (6-lane treadmill, UGO BASILE S.R.L, Italy) for 3 days: on the first day, mice were trained for 12 minutes running sessions, with gradually increasing speed to a maximum of 18 m/minutes at a 10° inclination. The second and the third days, mice were trained for 18 minutes running sessions with gradually increasing speed to a maximum of 24m/minutes at a 10° inclination. After 3 days of acclimatization to the treadmill exercise, exhaustion and fatigue tests were carried out. After 4 minutes of warm up at a speed of 5m/minutes with a slope fixed at 10° inclination, the speed was increased to 14 m/minutes for 2 minutes. The test consisted of a run at gradually increased speed to a maximum of 32 m/minutes: every 2 minutes the speed was increased by 2 m/minutes until animals were exhausted. Exhaustion was defined when mice were unable to maintain the pace and spent 10 consecutive seconds in the “fatigue zone.” Therefore, mice were considered exhausted when they remained in the lower part of the treadmill (last third) for 10 seconds. The running time was measured.

### Echocardiography

Left ventricular EF and LV dimension were measured on a high-resolution echocardiographic system equipped with a 30-MHz mechanical transducer (VEVO 2100, VisualSonics Inc) as previously described ^15,16^. Mice were anesthetized using 2% oxygenated isoflurane by inhalation. Mice were anchored to a warming platform in a supine position, limbs were taped to the echocardiograph electrodes, and chests were shaved and cleaned with a chemical hair remover to minimize ultrasound attenuation. UNI’GEL ECG (Asept Inmed), from which all air bubbles had been expelled, was applied to the thorax to optimize the visibility of the cardiac chambers. EF was evaluated by planimetry as recommended ^17^. Two-dimensional, parasternal long-axis, and short-axis views were acquired, and the endocardial area of each frame was calculated by tracing the endocardial limits in the long-axis view, then the minimal and maximal areas were used to determine the left ventricular end-systolic and end-diastolic volumes, respectively. The system software uses a formula based on a cylindrical- hemi ellipsoid model (volume=8×area2/3π/length). The left ventricular EF was derived from the following formula: (end-diastolic volume−end-systolic volume)/end-diastolic volume×100. The cardiac wall thickness, LV posterior wall, interventricular septum, and LV internal diameter were calculated by tracing wall limits in both the long and short axis views.

### LV Pressure/Systolic Blood Pressure Measurement

LV diastolic pressure measurement was assessed via invasive catheterization technique. Briefly, mice were anesthetized with 2% isoflurane. A Scisense pressure catheter (Transonic) was inserted into the LV through the common carotid artery. Pressure was recorded using LabChart software. End-diastolic pressure, dP/dt minimum and maximum, Tau, and heart rate were automatically calculated by a curve fit through end-systolic and end-diastolic points on the pressure plot.

### Blood sampling for biochemical marker analysis/NFS

Blood samples were collected by the heparin retroorbital bleeding method at sacrifice. Blood cell counts were determined using an automated counter (scil Vet abc Plus+). Plasma was separated by a 10-min centrifugation at 2500 g and then stored at −80°C. Concentrations of the following biomarkers were measured using an Architect CI8200 analyzer (Abbott Diagnostics, North Chicago, Illinois, USA): triglycerides, using the lipoprotein-lipase/glycerol kinase/oxidase/peroxidase method; total cholesterol, using the esterase/oxidase/peroxidase method; and HDL cholesterol, using the accelerator/selective detergent/esterase/oxidase/peroxidase method. LDL cholesterol was then estimated using the Friedewald formula (LDL cholesterol [mmol/L] = total cholesterol – HDL cholesterol – [triglycerides/2,2], or LDL cholesterol [mg/dL] = total cholesterol – HDL cholesterol – [triglycerides/5]).

### Lung and heart Edema Assessment

To assess water content, the whole lung and heart were weighed immediately after being harvested (wet weight) and then dried for 7 days in an oven at 60 °C (dry weight). Tissue edema was quantified by calculating percentage of water content.

### Immunostaining

Prior to harvest, heart was perfused with 8 g/L containing NaCl 0.9% to stop the heart in diastole, it was then fixed with methanol, paraffin-embedded and cut into 7 μm thick sections. Alternatively, heart was fresh frozen in OCT and then cut into 7 μm thick sections.

Capillary density was evaluated in sections stained with anti-CD31 or anti-Podocalyxin (PODXL) antibodies by quantifying the number of CD31+ or PODXL+ elements. Muscularized vessels were identified using anti–alpha-smooth muscle actin (SMA) antibodies. Pericyte were identified using anti-NG2 antibodies. Leucocyte, macrophage, T-cell and B-cell infiltrations were quantified using anti-CD45, anti-CD68, anti-CD3 and anti-B220 antibodies, respectively. Endothelial adherens junction was characterized using anti-CDH5 antibodies. Edema was measured after FGB staining of muscle sections and quantified as the mean FGB-positive areas. Endothelial activation was identified using anti-ICAM-1 and anti-VCAM-1 antibodies and quantified as the % of ICAM-1 and VCAM-1 positive area respectively. Thrombi were identified using anti-CD41 antibodies and quantified as the number of CD41+ elements. (See Supplemental table 1 for antibody references.)

Cardiomyocyte mean surface area was measured using ImageJ software after membrane staining with Wheat Germ Agglutinin (WGA), Alexa Fluor™ 488 Conjugate (Invitrogen). Fibrosis was assessed either after sirius red staining or type I collagen (COL1A1) staining of heart sections by quantifying the percentage of red stained area.

Quantifications were done on images were acquired using an Axiozoom V16 or Axioscope A1 (Zeiss) under 200x magnification. They were conducted on 10 randomly selected images were performed using ImageJ/Fiji v2.0.0-rc-59 software (National Institute of Health, USA) by an investigator blinded to genotype. More precisely, each sample received a unique number. At the end of the experiment, the genotype/treatment for each sample was unveiled to enable data analysis.

For immunohistochemical analyses, primary antibodies were sequentially coupled with biotin-conjugated secondary antibodies and streptavidin-horseradish peroxidase (HRP) complex (Amersham). Staining was then revealed with a DAB substrate kit (Vector Laboratories) and tissue sections were counterstained with hematoxylin (see Supplemental table 2 for secondary antibody references).

For immunofluorescence analyzes, primary antibodies were resolved with Alexa-Fluor–conjugated secondary polyclonal antibodies (see Supplemental table 2). Nuclei were counterstained with DAPI (4′,6-diamidino-2-phenylindole).

For both immunohistochemical and immunofluorescence analyses, negative control experiments to check for antibody specificity were performed using secondary antibodies.

Representative confocal images were taken with a ZEISS LSM 700 scanning confocal microscope (Carl Zeiss, Germany).

### Quantitative RT-PCR

RNA was isolated using Tri Reagent (Molecular Research Center Inc) as instructed by the manufacturer, from heart tissue that had been snap-frozen in liquid nitrogen and homogenized. For quantitative real time-polymerase chain reaction (PCR) analyses, total RNA was reverse transcribed with M-MLV reverse transcriptase (Promega), and amplification was performed on an AriaMx Real Time PCR system (Agilent Technologies) using B-R SYBER Green SuperMix (Quanta Biosciences). Primer sequences are reported in Supplemental table 3.

The relative expression of each mRNA was calculated by the comparative threshold cycle method and normalized to b-actin rRNA expression.

### Western Blot Analysis

ATP2A2 protein level was evaluated by SDS PAGE using rabbit anti-ATP2A2 antibodies (Badrilla, Cat# A010-80). PLN phosphorylation at Serine 16 and Threonin 17 was evaluated by SDS PAGE using rabbit anti-phospho-PLN Ser16 (Badrilla, Cat# A010-12), rabbit anti-phospho-PLN Thr17 (Badrilla, Cat# A010-13) and mouse anti-total PLN (Badrilla, Cat# A010-14). RYR2 phosphorylation was evaluated by SDS PAGE using rabbit anti-phosphoRYR2 antibodies (Badrilla, Cat# A010-31).

Protein loading quantity was controlled using mouse monoclonal anti-α-tubulin antibodies (Sigma, Cat# T5168).

### RNA Sequencing

RNA was isolated using Tri Reagent^®^ (Molecular Research Center Inc) as instructed by the manufacturer from the cardiac vascular fraction of Leprdb/db mice and control Leprdb/+ mice. mRNA library preparation were realized following manufacturer’s recommendations (KAPA mRNA HyperPrep Kit from ROCHE). Final samples pooled library prep were sequenced on Nextseq 500 ILLUMINA, corresponding to 2×30Millions of reads per sample after demultiplexing.

Quality of raw data has been evaluated with FastQC 16. Poor quality sequences has been trimmed or removed with Trimmomatic 17 software to retain only good quality paired reads. Star v2.5.3a 18has been used to align reads on mm10 reference genome using standard options. Quantification of gene and isoform abundances has been done with rsem 1.2.28, prior to normalisation with edgeR bioconductor package 19. Finally, differential analysis has been conducted with the glm framework likelihood ratio test from edgeR. Multiple hypothesis adjusted p-values were calculated with the Benjamini-Hochberg procedure to control FDR. The FPKM values of all transcripts of which expression was significantly different between both groups in Supplemental table 4.

### Statistics

Results are reported as mean±SEM. Comparisons between groups were analysed for significance with the nonparametric Mann-Whitney test or the Kruskal-Wallis test followed by Dunn multiple comparison test (for >2 groups) using GraphPad Prism v8.0.2 (GraphPad Inc, San Diego, Calif). The normality and variance were not tested. Differences between groups were considered significant when P≤0.05 (*P≤0.05; **P≤0.01; ***P≤0.001).

## RESULTS

### Ovariectomized female mice fed with a HFD+L-NAME regimen display increased end diastolic pressure with preserved ejection fraction

In order to fit better with the HFpEF patient’s population which includes a large proportion of menopaused women, we chose to submit ovariectomized female mice to the HFD + L-NAME regimen proposed by G. Schiattarella et al ^12^. More specifically C57BL/6 female were ovariectomized at 7 weeks of age, The HFD was initiated 1 week later and the L-NAME treatment 3 weeks later. Notably L-NAME (1 mg/L) was given in the drinking water discontinuously i.e. 4 days/week, to prevent mice to lose weight. Assessments were performed 12 weeks and 10 weeks after the HFD regimen and L-NAME treatment were initiated respectively.

Firstly, we checked whether or not these mice developed hypertension, dyslipidemia and systemic low-grade inflammation. Discontinuous L-NAME was not sufficient to induce significant elevation of blood pressure in OXX female mice (Supplemental Figure 1A-C). However, as expected, mice fed with HFD gained weight (Supplemental Figure 1D). This was associated with an elevation of total cholesterol, HDL-cholesterol and LDL-cholesterol levels while triglycerides were not modulated (Supplemental Figure 1E-H). Notably, HFD also induced low grade inflammation attested with significantly increased circulating leucocytes (notably monocytes and lymphocytes) especially in HFD+L-NAME treated mice (Supplemental Figure 1I-K).

We then assessed cardiac function in these mice essentially via echocardiography and LV catheterization. In the four groups left ventricular ejection fraction (LVEF) was normal and above 50% (Figure 1A). On the contrary, HFD+L-NAME-treated OVX female mice displayed diastolic dysfunction attested by significantly elevated left ventricular end diastolic pressure (LVEDP) and relaxation constant Tau (Figure 1B-C). Mice treated with HFD or L-NAME alone displayed intermediate phenotypes. Consistent with heart failure, HFD+L-NAME treated OVX female mice displayed exercise intolerance in the treadmill test (Figure 1D).

**Figure 1:**
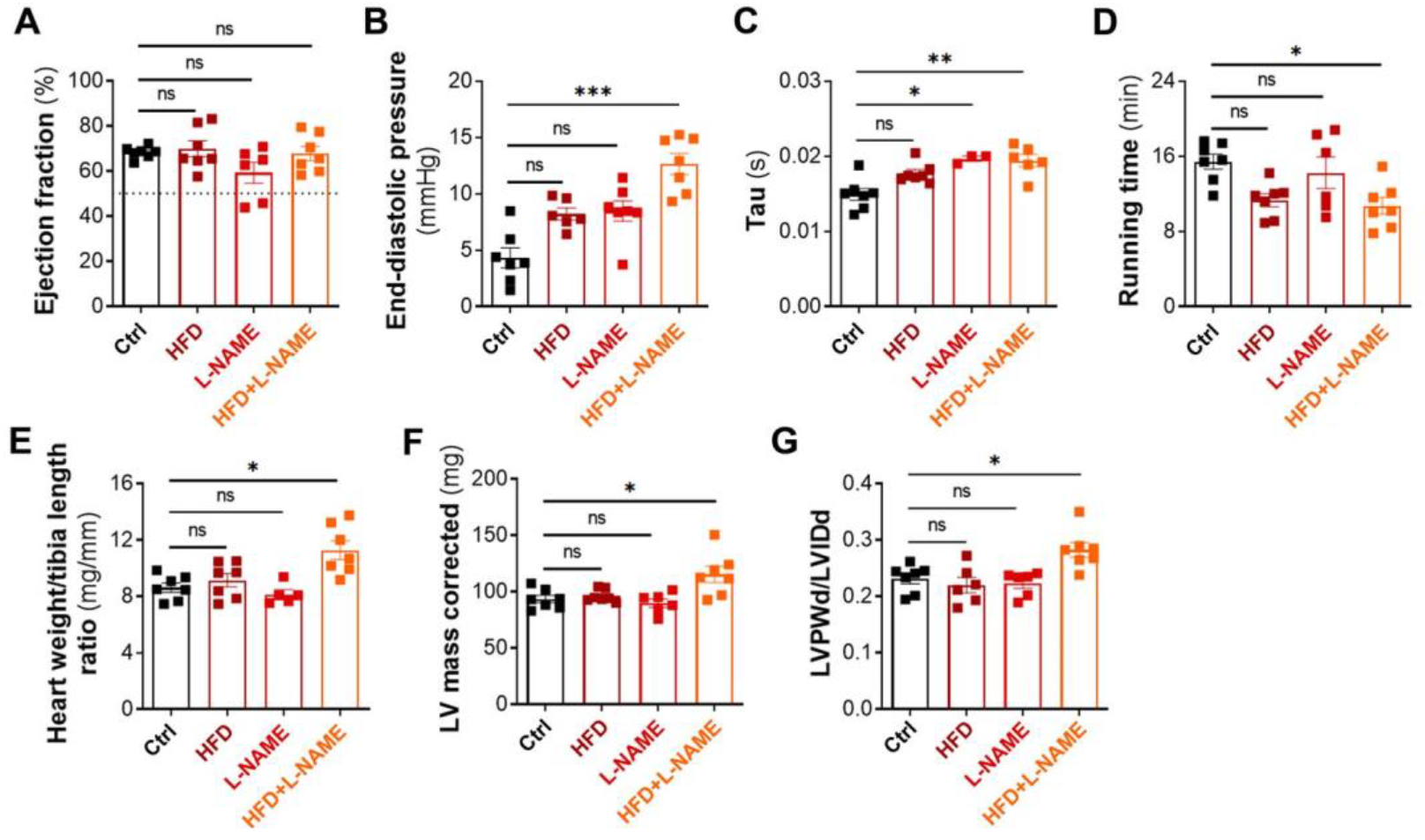
OVX female mice submitted to a HFD + L-NAME regimen develop diastolic dysfunction. C57BL/6 female mice were ovariectomized at 7 weeks of age, then submitted or not to a HFD regimen from 8 weeks of age and exposed or not to L-NAME (1 g/L in the drinking water) discontinuously (4 days/week) from 10 week of age. Cardiac structure and function was assessed at 20 weeks of age. (**A**) Ejection fraction was assessed via echocardiography (n=6-7). (**B**) LV end diastolic pressure (EDP) was measured by left ventricular catheterization (n=6-7). (**C**) Tau was measured by left ventricular catheterization (n=6-7). (**D**) Running time was assessed on treadmill (n=6-7). (E) The heart weight over tibia length was measured (n=5-7). (**F**) LV mass was assessed by echocardiography (n=6-7). (G) Diastolic left ventricular posterior wall (LVPWd) thickness and diastolic left ventricular internal diameter (LVIDd) were measured by echocardiography (n=6-7). Ratio of LVPWd and LVIDd was calculated. *: p≤0.05, **: p≤0.01, ***: p≤0.001, ns: not significant (Kruskal-Wallis test with Bonferroni post-hoc).

Assessment of heart weight/tibia length ratio showed HFD+L-NAME-treated OVX female mice developed cardiac hypertrophy (Figure 1E). This result was confirmed by echocardiographic measurements. Indeed both the LV mass and the diastolic LV posterior wall thickness (LVPWd)/LV internal diameter (LVIDd) ratio were significantly increased in HFD+L-NAME-treated OVX female mice (Figure 1F-G) confirming and attesting cardiac concentric hypertrophy. Neither the HFD regimen nor L-NAME treatment alone did induce cardiac hypertrophy.

In conclusion, consistent with Schiattarella et al. data, a HFD regimen combined with a L-NAME treatment synergistically induce cardiac hypertrophy and diastolic dysfunction in mice.

### Diastolic dysfunction observed in HFD+L-NAME-treated OVX female mice was associated with cardiomyocyte hypertrophy, fibrosis and inflammation

To further characterize cardiac remodelling associated with diastolic dysfunction in HFD+L-NAME-treated OVX female mice; we performed histological and gene expression analysis on the heart.

At cellular level, cardiac hypertrophy in HFD+L-NAME-fed OVX female mice was associated with increased cardiomyocyte size (Supplemental Figure 2A-B). Besides, *Myh7* mRNA expression signing cardiomyocyte dedifferentiation was not modified (Supplemental Figure 2C) while *Nos2* mRNA previously shown to be increased in this model and to participate in the development of diastolic dysfunction by Schiattarella et al. ^12^ was significantly increased (Supplemental Figure 2D).

Cardiomyocytes also showed altered Ca^2+^ homeostasis since we found HFD+L-NAME-fed OVX female had significantly decreased cardiac ATP2A2 protein levels (Supplemental Figure 2E,F). Phosphorylation of PLN on serine 16 was not affected while phosphorylation of PLN on threonine 17 tended to decreased (Supplemental Figure 2G-I). Phosphorylated RYR2 levels were not different between HFD+L-NAME-fed OVX female and control OVX female mice (Supplemental Figure 2J-K).

Notably, consistent with human HFpEF, diastolic dysfunction in HFD+L-NAME-fed OVX female mice was associated with increased perivascular and interstitial cardiac fibrosis (Supplemental Figure 3A-B). Fibrosis was found to be evaluated by both Sirius red and type I Collagen staining of heart sections and confirmed by qPCR analysis of *Col1a1* and *Col3a1* mRNA expression (Supplemental Figure 3D-E).

Also CD45+ leukocyte infiltration in the cardiac tissue of HFD+L-NAME-treated OVX female mice was significantly increased (Supplemental Figure 3F-G). Infiltrated leukocytes were mostly CD68+ macrophages but also CD3+ T-cells and B220+ B-cells and all of these 3 populations of cells were significantly increased in the heart of HFD+L-NAME-treated OVX female mice in comparison to control mice (Supplemental Figure 3H-J). Neither the HFD regimen nor L-NAME treatment alone did induce cardiac fibrosis or inflammation.

Consistent with human HFpEF, HFD+L-NAME-fed OVX female mice display cardiac hypertrophy, fibrosis and inflammation. Besides, HFD+L-NAME-fed OVX female mice display decreased ATP2A2 levels consistent with impaired cardiomyocyte active relaxation. Importantly, the HFD regimen needs to be associated with L-NAME or vice versa to induce cardiac remodeling in OVX female mice.

### Ovariectomized female mice fed with a HFD+L-NAME display cardiac small vessel disease

Since our model of HFpEF recapitulate most features of human HFpEF, we then phenotyped the cardiac microvasculature of HFD+L-NAME OVX female mice, essentially by doing histological analyses.

First, we quantified cardiac capillary density and found that capillary density was not modified in any of the groups (Figure 2A-B). However, we found that SMA+ arteriole mean diameter was significantly decreased in OVX mice treated either with L-NAME alone or HFD+L-NAME (Figure 2C-D) which is consistent with the vasoconstrictor effect of L-NAME. To assess vascular permeability, we measured fibrinogen and albumin extravasation and found that both HFD alone and HFD associated with L-NAME induced significant vascular leakage. Notably while L-NAME did not affect vascular permeability alone, it did exacerbate the effect of the HFD regimen (Figure 2E-G). Both L-NAME alone and L-NAME associated with HFD promoted endothelial cell activation attested by ICAM1 overexpression (Figure 2I-J). None of the groups showed increased small vessel thrombosis (Figure 2K).

**Figure 2:**
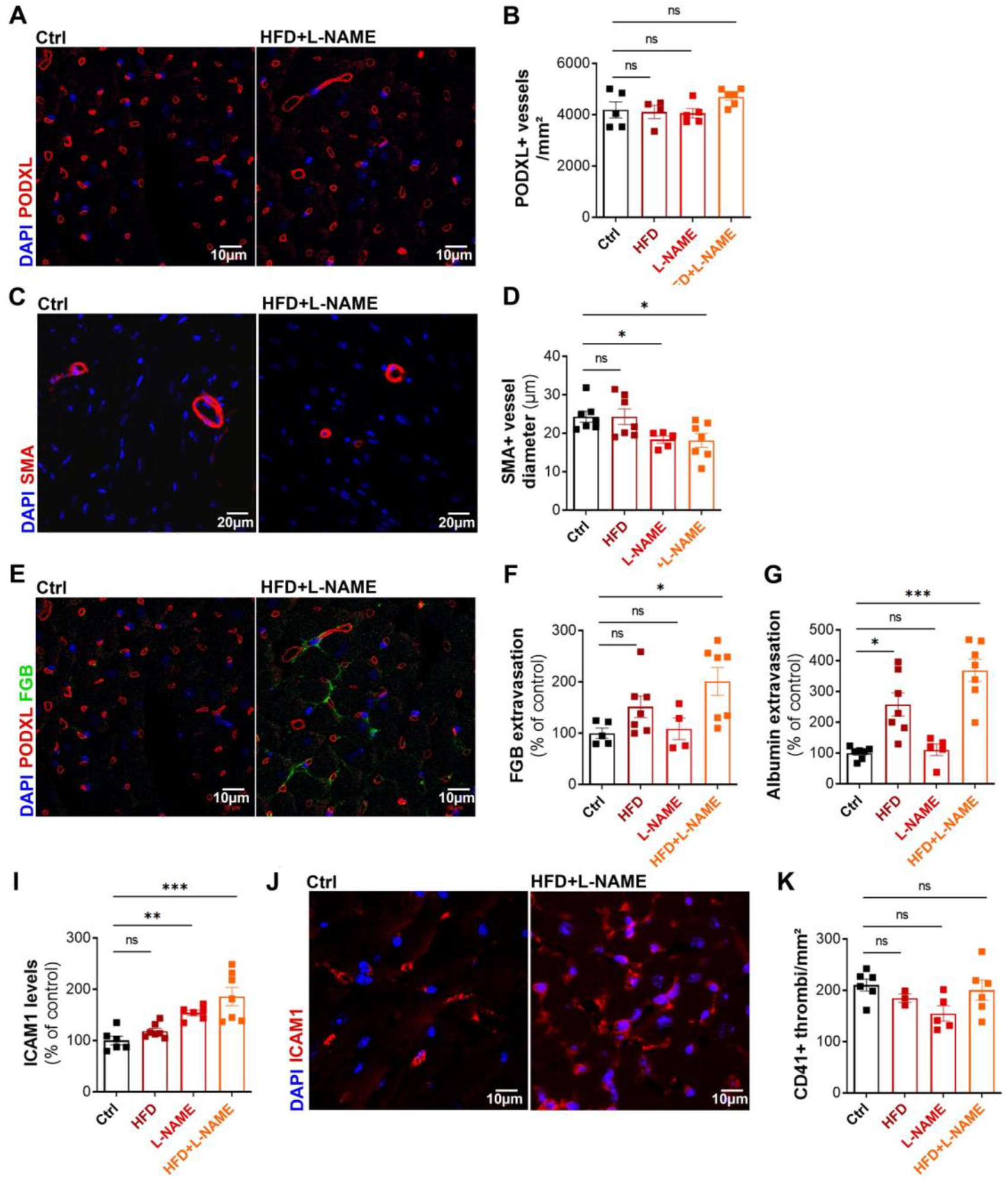
OVX female mice submitted to a HFD + L-NAME regimen display cardiac small vessel disease. C57BL/6 female mice were ovariectomized at 7 weeks of age, then submitted or not to a HFD regimen from 8 weeks of age and exposed or not to L-NAME (1 g/L in the drinking water) discontinuously (4 days/week) from 10 week of age. Mice were sacrificed at 20 weeks of age. (**A**) Heart cross sections were immunostained with anti-PODXL antibodies to identify ECs. (**B**) The number of PODXL+ capillary per mm^2^ was counted (n=5-7). (**C**) Heart cross sections were immuno-stained with anti-SMA (α smooth muscle actin) antibodies to identify smooth muscle cells. (**D**) The mean cardiac arteriole diameter was measured (n=5-7). (**E**) Heart cross sections were immunostained with anti-FGB antibodies. (**F**) FGB+ surface area was measured using Image J software (n=5-7). (**G**) Albumin+ surface area was measured using Image J software after immunostaining with anti-albumin antibodies (n=5-7). (**H**) ICAM1+ surface area was quantified using Image J software (n=6-7). (**I**) Heart cross sections were immunostained with anti-ICAM-1 antibodies. (**J**) The number of CD41+ thrombi per mm^2^ was counted (n=5-6). *: p≤0.05, **: p≤0.01, ***: p≤0.001, ns: not significant (Kruskal-Wallis test with Bonferroni post-hoc)

Altogether, these results show that diastolic dysfunction, in HFD+L-NAME-fed OVX female mice, is associated with endothelial dysfunction.

### While males develop diastolic dysfunction upon a HFD + L-NAME regimen, non OVX females do not

Considering the possible sexual dimorphism associated with HFpEF we tested whether the same phenotypes were also observed in male and none OVX female.

As expected, LVEF was not modified by the HFD+L-NAME treatment neither in male nor in non-OVX females (Supplemental Figure 4A). However, while male developed diastolic dysfunction attested by increased EDP and Tau, non OVX female did not (Supplemental Figure 4B, C) suggesting that female hormones may be protective against HFpEF. Consistently, HFD+L-NAME treated male did show impaired exercise tolerance while non OVX female did not (Supplemental Figure 4D).

Notably, on the contrary to OVX females, males did not develop neither cardiac nor cardiomyocyte hypertrophy upon the HFD+L-NAME treatment (Supplemental Figure 4E-F) while they show significantly higher Myh7 mRNA expression suggesting cardiomyocyte dedifferentiation (Supplemental Figure 4G).

Cardiac fibrosis (both interstitial and perivascular) was increased only in OVX females under HFD+L-NAME regimen but not in male or non OVX female mice (Supplemental Figure 5A-B). Consistently Col1a1 and Col3a1 mRNA were also overexpressed only in OVX female mice (Supplemental Figure 5C-D).

Besides, the HFD+L-NAME regimen induced increased leucocyte infiltration only in OVX females but not non OVX and OVX females and males (Supplemental figure 5E).

Altogether, these results show that the cardiac remodeling associated with diastolic dysfunction upon a HFD+L-NAME regimen is different in males and females. In males, diastolic dysfunction was associated with increased Myh7 expression but not with cardiac hypertrophy, fibrosis or inflammation, while in OVX female diastolic dysfunction was associated with cardiac hypertrophy, and fibrosis. Notably cardiac inflammation was not associated with diastolic dysfunction since both OVX and non OVX female did show increased cardiac leukocyte infiltration while only OVX female did develop diastolic dysfunction.

### Males fed with a HDF + L-NAME regimen do not show cardiac small vessel disease

Then we assessed the phenotype of the cardiac microvasculature of male and non OVX females treated with HFD+L-NAME.

Capillary density was not modified by the HFD+L-NAME regimen in any of the groups (Figure 3A). However, the HFD+L-NAME treatment decreased the mean arteriole diameter in both OVX and non OVX female mice but not in males (Figure 3B). Similarly, FGB extravasation was increased in both OVX and non OVX females but not in males (Figure 3C) so was endothelial cell activation attested by increased *Sele* mRNA, and *Icam1* mRNA and protein expression (Figure 3D-F).

**Figure 3:**
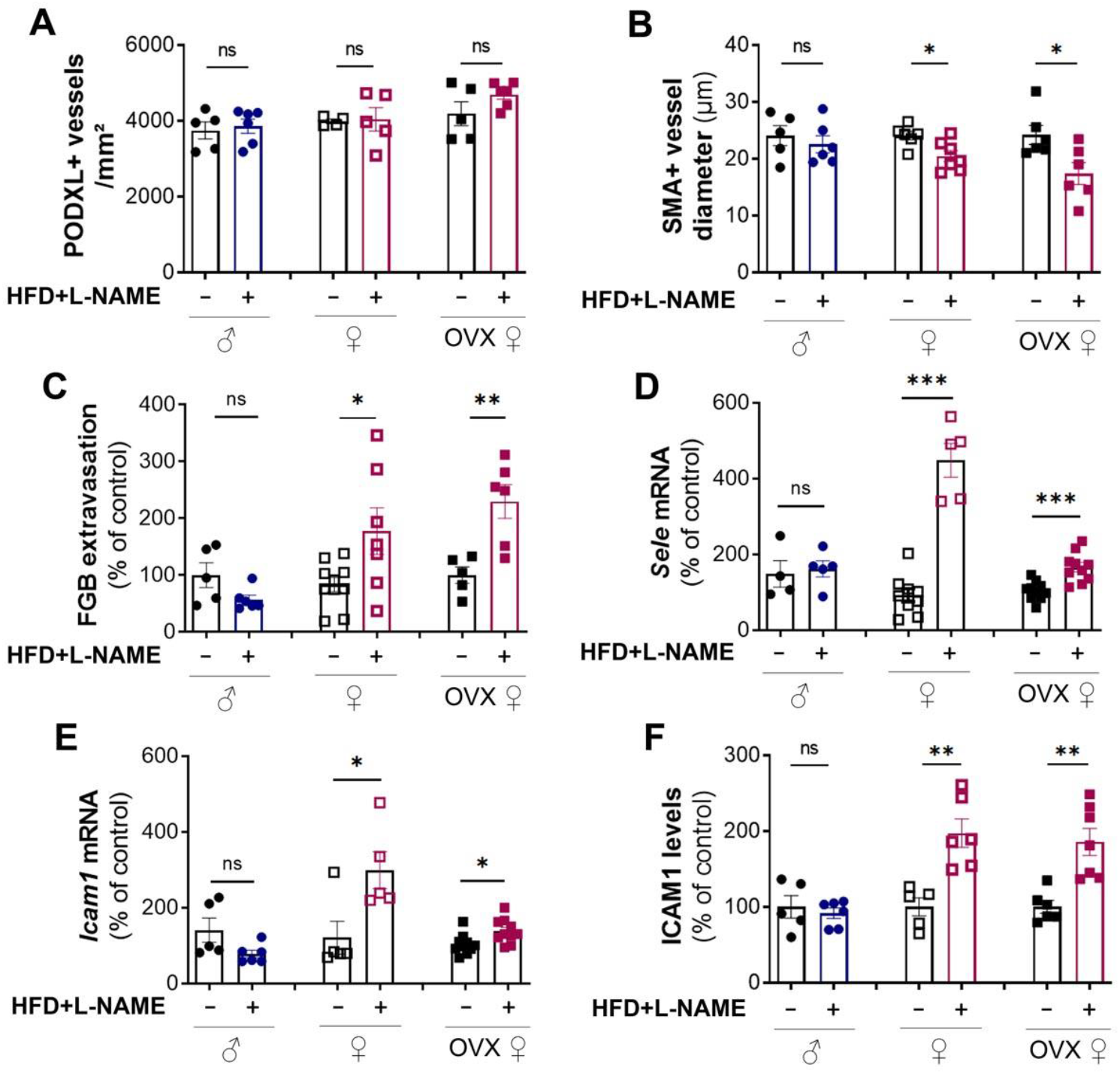
Only female mice submitted to a HFD + L-NAME regimen display cardiac small vessel disease. C57BL/6 male and female mice were ovariectomized or not at 7 weeks of age, then submitted or not to a HFD regimen from 8 weeks of age and exposed or not to L-NAME (1 g/L in the drinking water) discontinuously (4 days/week) from 10 week of age. Mice were sacrificed at 20 weeks of age. (**A**) Heart cross sections were immunostained with anti-PODXL antibodies to identify ECs. The number of PODXL+ capillary per mm^2^ was counted (n=5). (**B**) Heart cross sections were immuno-stained with anti-SMA antibodies to identify smooth muscle cells. The mean cardiac arteriole diameter was measured (n=5-7). (**C**) Heart cross sections were immunostained with anti-FGB antibodies. FGB+ surface area was measured using Image J software (n=5-7). (**D**) Sele mRNA expression was quantified by RT-qPCR in total heart extract and normalized to Actb mRNA (n=4-8). Icam1 mRNA expression was quantified by RT-qPCR in total heart extract and normalized to Actb mRNA (n=5-8). (**F**) ICAM1+ surface area was quantified using Image J software (n=5-7). *: p≤0.05, **: p≤0.01, ***: p≤0.001, ns: not significant (Mann-Whitney test).

Altogether these results demonstrate that cardiac small vessel disease only develops in female mice upon the HFD+L-NAME regimen. In order to investigate whether or not cardiac small vessel disease may participate in the development of diastolic dysfunction in OVX female mice, we used a mouse model in which we have previously shown endothelial integrity is preserved upon stress ^13^ i.e. endothelial specific *Cdon* deficient mice.

### Cdon endothelial KO protects ECs in OVX female mice submitted to the HFD+L-NAME treatment

We first verified these mice are protected from endothelial dysfunction when fed with the HFD + L-NAME regimen. *Cdon* endothelial KO (*Cdon*^ECKO^) did not modify microvessel density (Figure 4A-B) upon the HFD+L-NAME regimen. However, it did increase arteriole diameter (Figure 4C-D) and prevent HFD+L-NAME-induced abnormal endothelium permeability and activation, attested by decreased fibrinogen extravasation (Figure 4E-F) and decreased ICAM1 levels respectively (Figure 4G-H). Consistent with the decreased endothelium activation, CD45+ leukocyte, especially CD68+ macrophage, recruitment was significantly decreased in OVX female *Cdon*^ECKO^ mice in comparison to control mice treated with HFD+L-NAME (Figure 4I-K).

**Figure 4:**
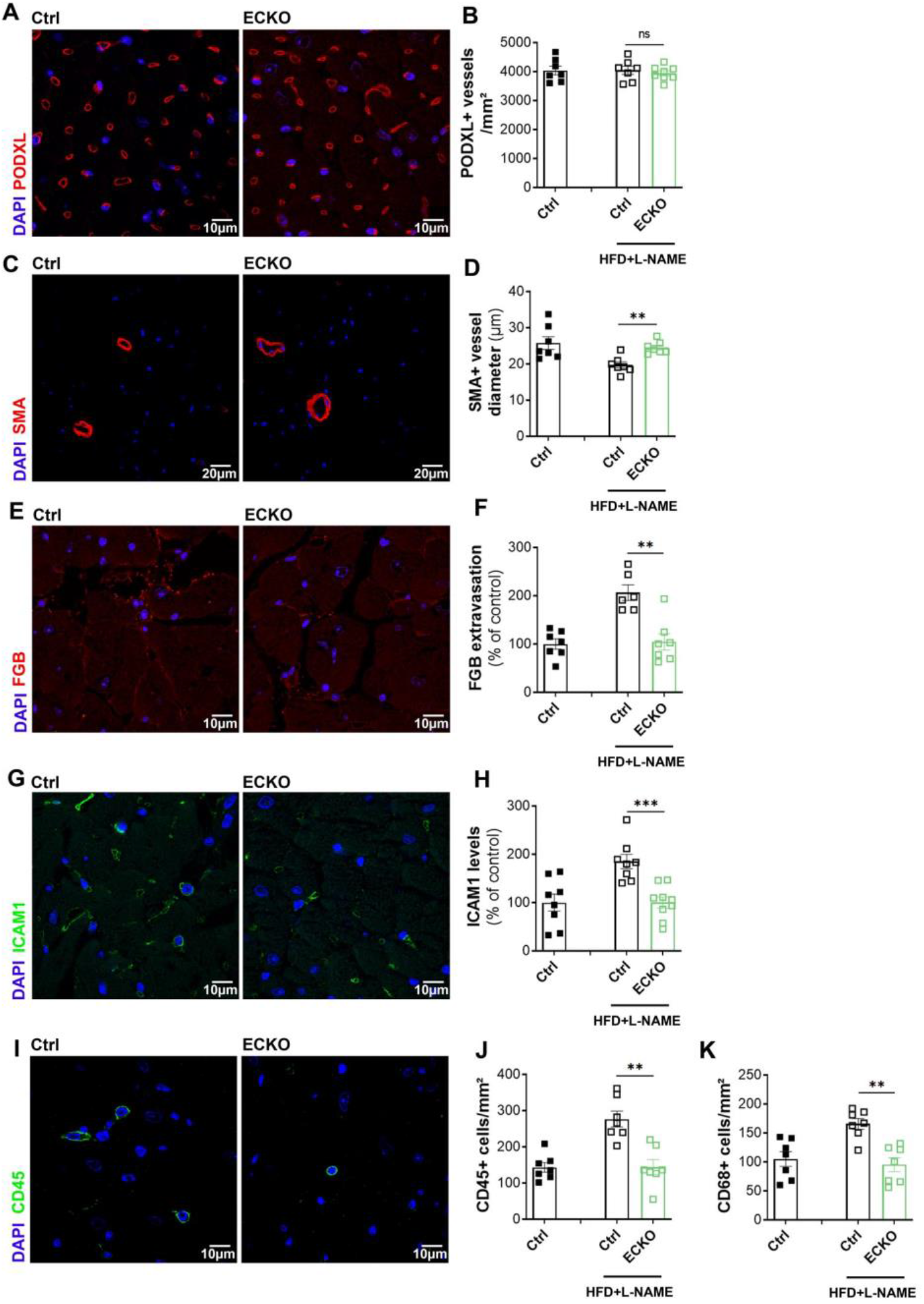
OVX Cdon^ECKO^ female mice submitted to a HFD + L-NAME regimen are protected from cardiac small vessel disease. Cdh5-Cre/ERT2; Cdon^Flox/Flox^ (Cdon^ECKO^) female mice and their control littermate (Cdon^Flox/Flox^) were ovariectomized or not at 7 weeks of age, then submitted or not to a HFD regimen from 8 weeks of age and exposed or not to L-NAME (1 g/L in the drinking water) discontinuously (4 days/week) from 10 week of age. Mice were sacrificed at 20 weeks of age. (**A**) Heart cross sections were immunostained with anti-PODXL antibodies to identify ECs. (**B**) The number of PODXL+ capillary per mm^2^ was counted (n=7). (**C**) Heart cross sections were immuno-stained with anti-SMA (α smooth muscle actin) antibodies to identify smooth muscle cells. (**D**) The mean cardiac arteriole diameter was measured (n=7). (**E**) Heart cross sections were immunostained with anti-FGB antibodies. (**F**) FGB+ surface area was measured using Image J software (n=6-7). (**G**) Heart cross sections were immunostained with anti-ICAM-1 antibodies. (**H**) ICAM1+ surface area was quantified using Image J software (n=6-7). (**I**) Heart cross sections were immunostained with anti-CD45 antibodies to identify leukocytes. (**J**) The number of CD45+ leukocytes per mm^2^ was counted (n=7). (**K**) The number of CD68+ monocytes/macrophages per mm^2^ was counted (n=7). *: p≤0.05, **: p≤0.01, ***: p≤0.001, ns: not significant (Mann-Whitney test)

These results showed that *Cdon* endothelial KO protects cardiac ECs and prevents cardiac inflammation in OVX female mice fed with a HFD+L-NAME regimen indicating these mice can be used to investigate the role of endothelial dysfunction in the pathophysiology of diastolic dysfunction.

### Improving cardiac small vessel disease does not prevent the occurrence of diastolic dysfunction in OVX female mice

Then we tested whether or not these mice develop diastolic dysfunction. As expected, systolic function, attested by a LVEF above 50%, was normal in OVX *Cdon*^ECKO^ female mice treated with the HFD+L-NAME regimen (Figure 5A). Importantly, the HFD+L-NAME regimen increased LVEDP and tau in both OVX *Cdon*^WT^ and OVX *Cdon*^ECKO^ female mice (Figure 5B-C) meaning that *Cdon*^ECKO^ mice develop diastolic dysfunction dysfunction in the absence of endothelial dysfunction and cardiac inflammation. Consistently, OVX *Cdon*^ECKO^ female mice fed with the HFD+L-NAME regimen display exercise intolerance just like *Cdon*^WT^ mice (Figure 5D).

**Figure 5:**
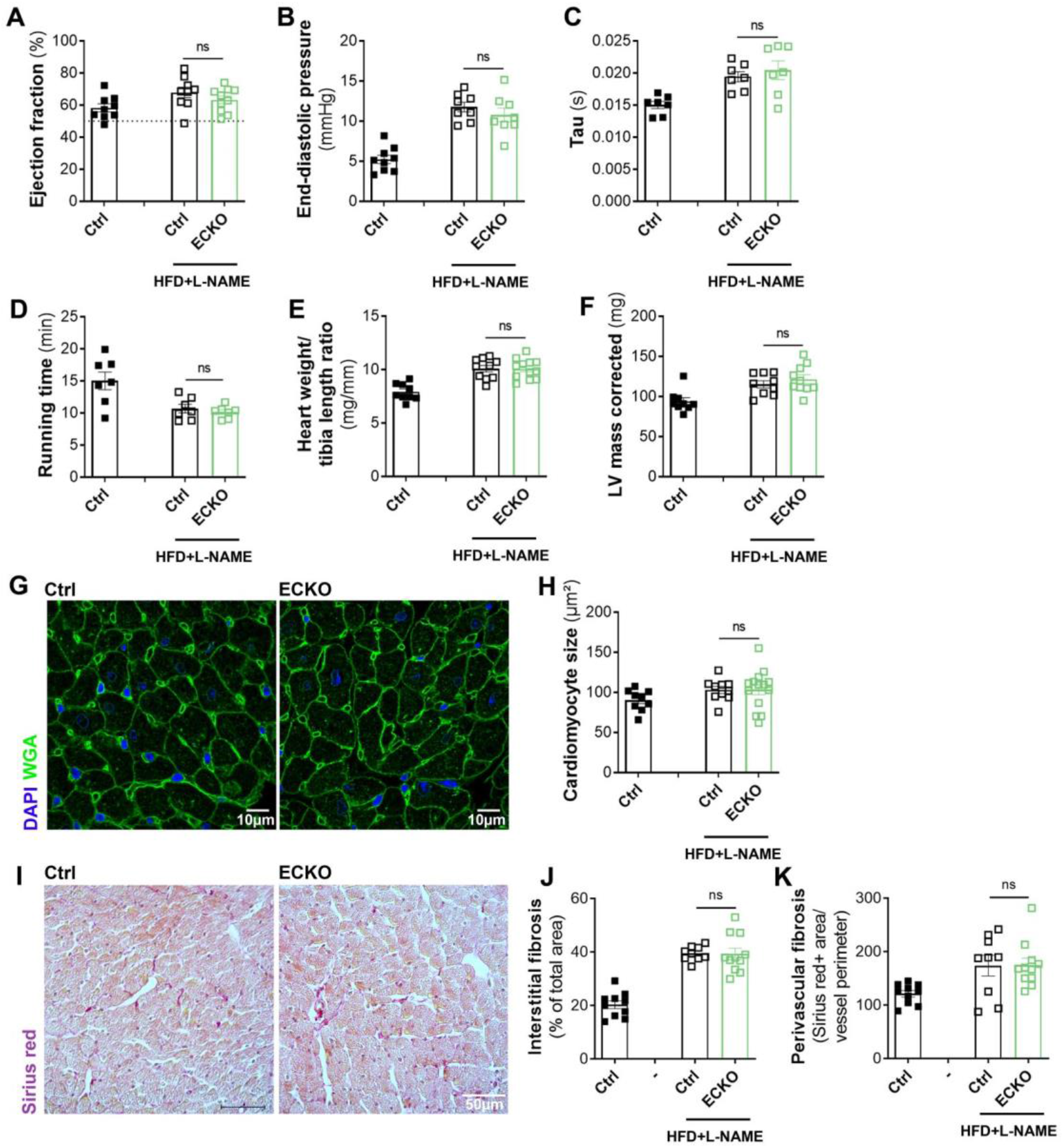
OVX Cdon^ECKO^ female mice submitted to a HFD + L-NAME regimen develop diastolic dysfunction. Cdh5-Cre/ERT2; Cdon^Flox/Flox^ (Cdon^ECKO^) female mice and their control littermate (Cdon^Flox/Flox^) were ovariectomized or not at 7 weeks of age, then submitted or not to a HFD regimen from 8 weeks of age and exposed or not to L-NAME (1 g/L in the drinking water) discontinuously (4 days/week) from 10 week of age. Mice were sacrificed at 20 weeks of age. (**A**) Ejection fraction was assessed via echocardiography (n=9). (**B**) LV end diastolic pressure (EDP) was measured by left ventricular catheterization (n=9). (**C**) Tau was measured by left ventricular catheterization (n=7). (**D**) Running time was assessed on treadmill (n=7). (**E**) The heart weight over tibia length was measured (n=9-12). LV mass was assessed by echocardiography (n=9). (**G**) Heart cross sections were stained with FITC-labelled WGA. (**H**) Cardiomyocyte cross section area was measured using Image J software (n=9-12). (**I**) Heart cross sections were stained with Sirius red to identify fibrosis. (**J**) Interstitial fibrosis was quantified using Image J software (n=10). (**K**) Perivascular fibrosis was quantified using Image J software (n=9). *: p≤0.05, **: p≤0.01, ***: p≤0.001, ns: not significant (Mann-Whitney test)

Notably, OVX *Cdon*^ECKO^ female mice develop cardiac hypertrophy and fibrosis just like control mice. More specifically, Cardiac mass, measured either with a scale or via echocardiography was identically in OVX *Cdon*^ECKO^ mice and OVX *Cdon*^WT^ mice treated with the HFD+L-NAME regimen (Figure 5E-F) so was cardiomyocyte size measured after WGA staining (Figure 5G-H). Similarly, Sirius red staining showed that both interstitial and perivascular fibrosis was identical in OVX *Cdon*^ECKO^ mice and OVX *Cdon*^WT^ mice treated with the HFD+L-NAME regimen (Figure 5I-J)

Altogether, these results reveal that OVX *Cdon*^ECKO^ female mice fed with and HDF+L-NAME regimen develop cardiac fibrosis, hypertrophy and diastolic dysfunction even though they do not display increased micro-vessel permeability and endothelial activation suggesting that micro-vessel permeability and activation are not the unique triggers of diastolic dysfunction in OVX female mice fed with an HFD + L-NAME regimen.

### Colchicine therapy *does not prevent the development of diastolic dysfunction in OVX female mice*

Notably OVX Cdon^ECKO^ female mice displayed reduced cardiac inflammation upon HFD+L-NAME regimen suggesting that increased leucocyte infiltration in the heart may also not promote the development of diastolic dysfunction. To test this hypothesis, OVX female mice were treated or not with colchicine, an anti-inflammatory drug widely used in cardiology, for 10 weeks. Colchicine treatment was initiated 2 weeks after the HFD regimen at the same time as the L-NAME treatment.

As expected colchicine treatment reduced both circulating and cardiac infiltrated leucocytes (Figure 6A-C). Cardiac dimensions and function were then assessed via echocardiography and LV catheterization. LVEF was not modified by the colchicine treatment neither were LVEDP and Tau in OVX female mice treated with the HFD+L-NAME regimen (Figure 6D-F) suggesting that cardiac infiltration with leucocyte is not necessary for diastolic dysfunction to appear. Consistently, colchicine treatment did not improve exercise tolerance in OVX female mice treated with the HFD+L-NAME regimen (Figure 6G).

**Figure 6.**
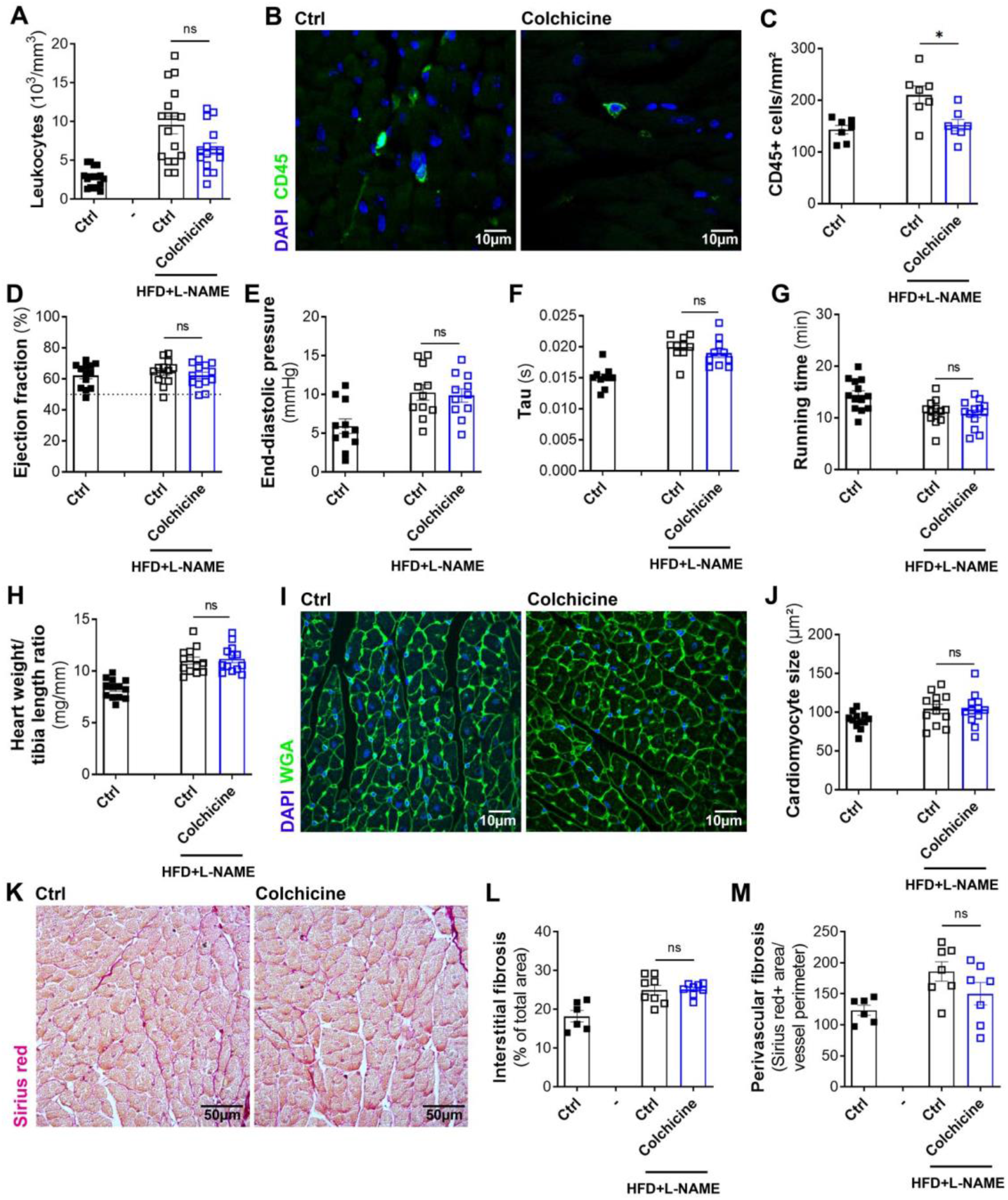
Colchicine therapy does not prevent the occurrence of diastolic dysfunction in OVX female mice submitted to a HFD + L-NAME regimen. C57BL/6 male and female mice were ovariectomized or not at 7 weeks of age, then submitted or not to a HFD regimen from 8 weeks of age and exposed or not to L-NAME (1 g/L in the drinking water) discontinuously (4 days/week) from 10 week of age. Colchicine treatment (200 μg/kg/day) was initiated from 10 weeks of age. Mice were sacrificed at 20 weeks of age. (**A**) Leukocyte count was calculated in total blood samples (n=10-17). (**B**) Heart cross sections were immunostained with anti-CD45 antibodies to identify leukocytes. (**C**) The number of CD45+ leukocytes per mm^2^ was counted (n=7). (**D**) Ejection fraction was assessed via echocardiography (n=10-17). (**E**) LV end diastolic pressure (EDP) was measured by left ventricular catheterization (n=11). (**F**) Tau was measured by left ventricular catheterization (n=10). (**G**) Running time was assessed on treadmill (n=13-17). (**H**) The heart weight over tibia length was measured (n=13-17). (**I**) Heart cross sections were stained with FITC-labelled WGA. (**J**) Cardiomyocyte cross section area was measured using Image J software (n=12). (**K**) Heart cross sections were stained with Sirius red to identify fibrosis. (**L**) Interstitial fibrosis was quantified using Image J software (n=6-8). (**M**) Perivascular fibrosis was quantified using Image J software (n=6-8). *: p≤0.05, **: p≤0.01, ***: p≤0.001, ns: not significant (Mann-Whitney test)

Notably, colchicine administration did not modify cardiac hypertrophy (Figure 6H-J) and fibrosis (Figure 6K-M) either.

Altogether these data confirm and demonstrate that increased leucocyte infiltration in the heart does not promote diastolic dysfunction in the mouse model of HFpEF triggered by a HFD+L-NAME combination treatment.

### Nos2 overexpression is a common feature of diastolic dysfunction observed in male and OVX females fed with the HFD + L-NAME regimen

Since Schiattarella at al. previously demonstrated diastolic dysfunction was driven by nitrosative stress in the mouse model of HFpEF triggered by a HFD+L-NAME combination treatment. We tested diastolic dysfunction was associated with increased NOS2 expression in both male and OVX female. As shown Supplemental Figure 6, Nos2 mRNA was increased in the heart of both male and OVX female exposed to HFD + L-NAME, but not in the heart of none OVX female, supporting the critical role nitrostative stress in the pathophysiology of HFpEF.

## DISCUSSION

Endothelial dysfunction has been proposed to be causing HFpEF since almost 10 years in the paradigm proposed by Paulus et al in 2013 ^5^. The present study demonstrates for the first time that neither endothelium activation, nor endothelium leakage do participate in the pathophysiology of this disease at least in the mouse model of HFpEF triggered by a HFD + L-NAME regimen. Indeed, we found that diastolic dysfunction still develops in OVX female mice fed by a HFD + L-NAME regimen and “protected” from endothelium leakage and activation. Interestingly, this is not the first time endothelial dysfunction is suggested not to have a critical role in the pathophysiology of HFpEF since several therapies aiming at restoring the NO/cGMP pathway, another important feature of endothelium dysfunction failed at improving HFpEF patient conditions ^20^.

Importantly the causal role of endothelial dysfunction in the pathophysiology of HFpEF has not been yet excluded since endothelial dysfunction may affect cardiomyocyte biology and cardiac function in many ways ^11^. Notably, ECs are essential cell components of blood vessels which are necessary to bring nutriments and oxygen to cardiomyocytes. Endothelial dysfunction may then cause cardiac hypoxia and subsequent cardiomyocyte dysfunction. OVX female mice fed with and HFD + L-NAME regimen did not show capillary rarefaction or increased capillary thrombosis suggesting capillary perfusion is not altered in these mice; however the mean arteriole diameter was diminished. The importance of cardiac hypoxia remains to be identified. Besides, ECs were shown to produce signals necessary for cardiomyocyte homeostasis including NO. Whether or not modifications of such signals may participate in the pathophysiology of HFpEF remain to be tested. Finally, EC dysfunction may affect cardiomyocyte function indirectly by modifying cardiomyocyte microenvironment. Notably, EC dysfunction may promote cardiac fibrosis, cardiac inflammation or cardiac edema via increased production of pro-fibrotic, pro-inflammatory signals or impaired barrier properties. Our data indicate that neither cardiac inflammation nor cardiac edema seems to promote diastolic dysfunction. However, cardiac fibrosis which did develop in OVX Cdon^ECKO^ female mice fed with the HFD + L-NAME regimen may results from endothelial dysfunction ^21^ and promotes HFpEF.

Besides, in the present study, we have further explored the possible sexual dimorphism associated with the pathophysiology of HFpEF. Notably, we found that cardiac remodeling associated with HFpEF differs in males and females. While diastolic dysfunction was associated with cardiomyocyte dedifferentiation attested with increased Myh7 mRNA expression in male. The HFD + L-NAME regimen induced endothelial dysfunction and cardiac inflammation only in females. Cardiomyocyte hypertrophy and fibrosis was specifically seen in OVX females but not in non OVX females or males. Importantly, these data seem to be relevant to humans, since women show higher inflammatory biomarkers including C-reactive protein (CRP) in several large-scale proteomic studies ^23^. Besides, women were reported to display more concentric cardiac hypertrophy while men display more eccentric remodeling ^22^. Also, pre-menopausal women are better protected against cardiac hypertrophy compared with men, but this protection was shown to be abolished following menopause and is partially restored after estrogen replacement therapy ^26^. Finally, women with HFpEF were shown to have significantly higher plasma levels of the propeptide for type I collagen than men with HFpEF ^25^. Moreover, women with hypertrophic cardiomyopathy were reported to display more fibrosis than male patients ^24^. Notably, estradiol was shown to diminish Col1a1 and Col3a1 in cultured-fibroblasts ^27^ possibly explaining why we only observed cardiac fibrosis in the heart of OVX females.

Consistent with Schiattarella et al. data, we found that diastolic dysfunction induced by a HFD + L-NAME regimen was associated with increased NOS2 levels ^12^. This result was observed in both males and OVX females (Supplemental Figure 6). Whether or not, HFpEF is induced by common or specific mechanisms, in males and females, remains then to be determined. So far, the increased NOS2 levels are the only feature that is correlated with the presence of diastolic dysfunction.

In conclusion, the present paper reveals for the first time that neither endothelium activation, nor endothelium leakage participates in the pathophysiology of HFpEF at least in the mouse model of HFpEF triggered by a HFD + L-NAME regimen. These results together with the negative results of clinical trials targeting the NO/cGMP pathway weakens Paulus et al paradigm ^5^ and suggest that neither endothelial dysfunction nor cardiac inflammation is causing HFpEF. However, considering HFpEF is an heterogeneous disease with multiple pheno-groups ^28^, these results needs to be verified in other animal models of HFpEF. Indeed, even though endothelial dysfunction does not seem to be necessary for diastolic dysfunction to develop upon a HFD + L-NAME regimen, EC protection was previously shown to prevent the occurrence of both diastolic and systolic dysfunction in other setting including pressure overload and hypertension ^11^. Besides, modification of some EC properties was shown to be sufficient to induced diastolic dysfunction in the absence of any other cardiovascular risk factor ^6,8–10^. Similarly while limiting cardiac inflammation with colchicine does not prevent the occurrence of HFpEF in HFD + L-NAME fed OVX female mice, we previously found that mast cell stabilization was able to prevent the occurrence of diastolic dysfunction in diabetic obese Lepr^db/db^ mice ^29^.

Finally this paper further highlight the importance of considering sex differences when investigating the pathophysiology of HFpEF.

## Acknowledgments

We thank Annabel Reynaud, Sylvain Grolleau, and Maxime David for their technical help.

This work benefited from equipment and services from the iGenSeq (RNA sequencing) and iCONICS (RNAseq analysis) core facilities at the ICM (Institut du Cerveau et de la Moelle épinière, Hôpital Pitié-Salpêtrière, PARIS, France).

