## Supplemental data for "Increased endothelium activation and leakage do not promote diastolic dysfunction in mice fed with a high fat diet and treated with L-NAME"

### Supplementary data

**Supplemental Tables**

A/ List of primary antibodies

| Target antigen | Vendor or Source | Catalog # | Working concentration |
|---|---|---|---|
| mouse PECAM-1 (CD31) | BMA Biomedicals | T-2001 | 2 µg/mL |
| mouse PODXL | R&D systems | AF1556 | 4 µg/mL |
| mouse Cy3-anti-smα-actin | Sigma | C6198 | 5 µg/mL |
| mouse ACTN2 | Sigma | A 2172 | |
| mouse ICAM-1 (CD54) | BD Pharmingen | 550287 | 2.5 µg/mL |
| mouse VCAM-1 | Invitrogen | 14-1061-82 | |
| mouse FGB | Abcam | ab227063 | |
| mouse COL1A1 | Abcam | ab21285 | |
| mouse CD41 | Proteintech | 18308-1-AP | 3.5 µg/mL |
| mouse Myh7 | Atlas antibodies | HPA001239 | 1 µg/mL |
| mouse CD45 | BD Pharmingen | 550539 | 0.625 µg/mL |
| mouse CD68 | Biolegend | 137001 | 5 µg/mL |
| mouse CD3 | Santa-Cruz Biotechnology | sc-1127 | |
| mouse B220 | R&D systems | MAB1217 | |
| mouse ATP2A2 | Badrilla | A010-80 | |
| phospho-PLN (Ser16) | Badrilla | A010-12 | |
| phospho-PLN (Thr17) | Badrilla | A010-13 | |
| total-PLN | Badrilla | A010-14 | |
| phospho-RYR2 (Ser2814) | Badrilla | A010-31 | |
| α-tubulin | Sigma | T5168 | |

B/ List of secondary antibodies

| Target antigen | Conjugate | Vendor or Source | Catalog # | Working concentration |
|---|---|---|---|---|
| Goat IgG | Alexa Fluor 568 | Invitrogen | A-11057 | 10 µg/mL |
| Rat IgG | Alexa Fluor 647 | Invitrogen | A-48265 | 10 µg/mL |
| Hamster IgG | Biotin | Jackson ImmunoResearch | 127-065-160 | Dilution 1/500 |
| Goat IgG | Alexa Fluor 488 | Invitrogen | A-11055 | 10 µg/mL |
| Rabbit IgG | Alexa Fluor 488 | Invitrogen | A-21206 | 10 µg/mL |
| Rabbit IgG | Alexa Fluor 568 | Invitrogen | A-10042 | 10 µg/mL |

**Supplemental Table 1: List of antibodies used for immunostainings**

| | | |
|---|---|---|
| mouse Actb | F | 5′- -3′ |
| | R | 5′- -3′ |
| mouse Col1a1 | F | 5′-CAACCTCAAGAAGGCCCTGC-3′ |
| | R | 5′-TGTCCAAGGGAGCCACATCG-3′ |
| mouse Col3a1 | F | 5′-AGCACGAGGTCTTGCTGGAC-3′ |
| | R | 5′-ACCAGCTGTACCAGGCTGAC-3′ |
| mouse Myh7 | F | 5′-GGATGACGTCACCTCCAACA-3′ |
| | R | 5′-AGATCAGAGCCTCCTTCTCGT-3′ |
| mouse Ttn N-2B | F | 5′-ACAGTGGGAAAGCAAAGACATC-3′ |
| | R | 5′-AGGTGGCCCAGAGCTACTTC-3′ |
| mouse Ttn N2BA | F | 5′-GAGACATTGCTCCGCTTTTC-3′ |
| | R | 5′-GATCTCCAAAGAGGCTGTC-3′ |
| mouse Atp2a2 (Serca2a) | F | 5′-GATCCTCTACGTGGAACCTTTG-3′ |
| | R | 5′-GGTAGATGTGTTGCTAACAACG-3′ |

**Supplemental Table 2: List of primers used for reverse transcription (RT) quantitative polymer chain reaction (qPCR).** F: forward; R: reverse

**Supplemental Figures and figure legends**

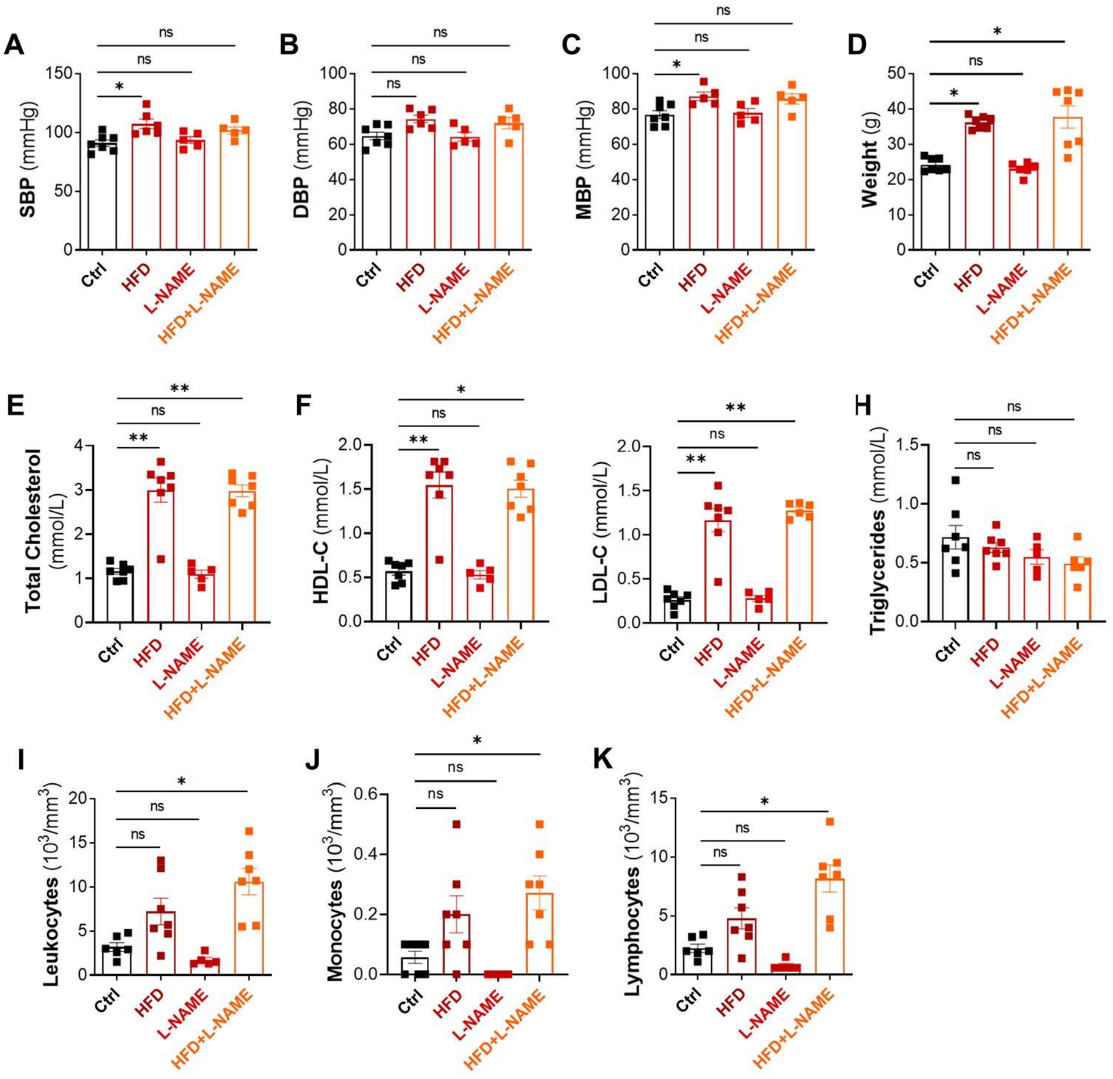

**Supplemental figure 1: OVX female mice submitted to a HFD + L-NAME regimen display overweight, dyslipidaemia and low grade inflammation.**

C57BL/6 female mice were ovariectomized at 7 weeks of age, then submitted or not to a HFD regimen from 8 weeks of age and exposed or not to L-NAME (1 g/L in the drinking water) discontinuously (4 days/week) from 10 week of age. Measurements were done at 20 weeks of age. (**A**) Systolic, (**B**) diastolic and (**C**) mean blood pressures were measured via left ventricular catheterization. (**D**) Weight of mice was measure in each group. (**E**) Total cholesterol, (**F**) HDL-cholesterol, (**G**), LDL-cholesterol and (**H**) triglycerides, were measured in plasma samples. (**I**) Leukocytes count, (**J**) monocyte count and (**K**) lymphocyte count were calculated in total blood samples.

*: p≤0.05, **: p≤0.01, ***: p≤0.001, ns: not significant (Kruskal-Wallis test with Bonferroni post hoc)

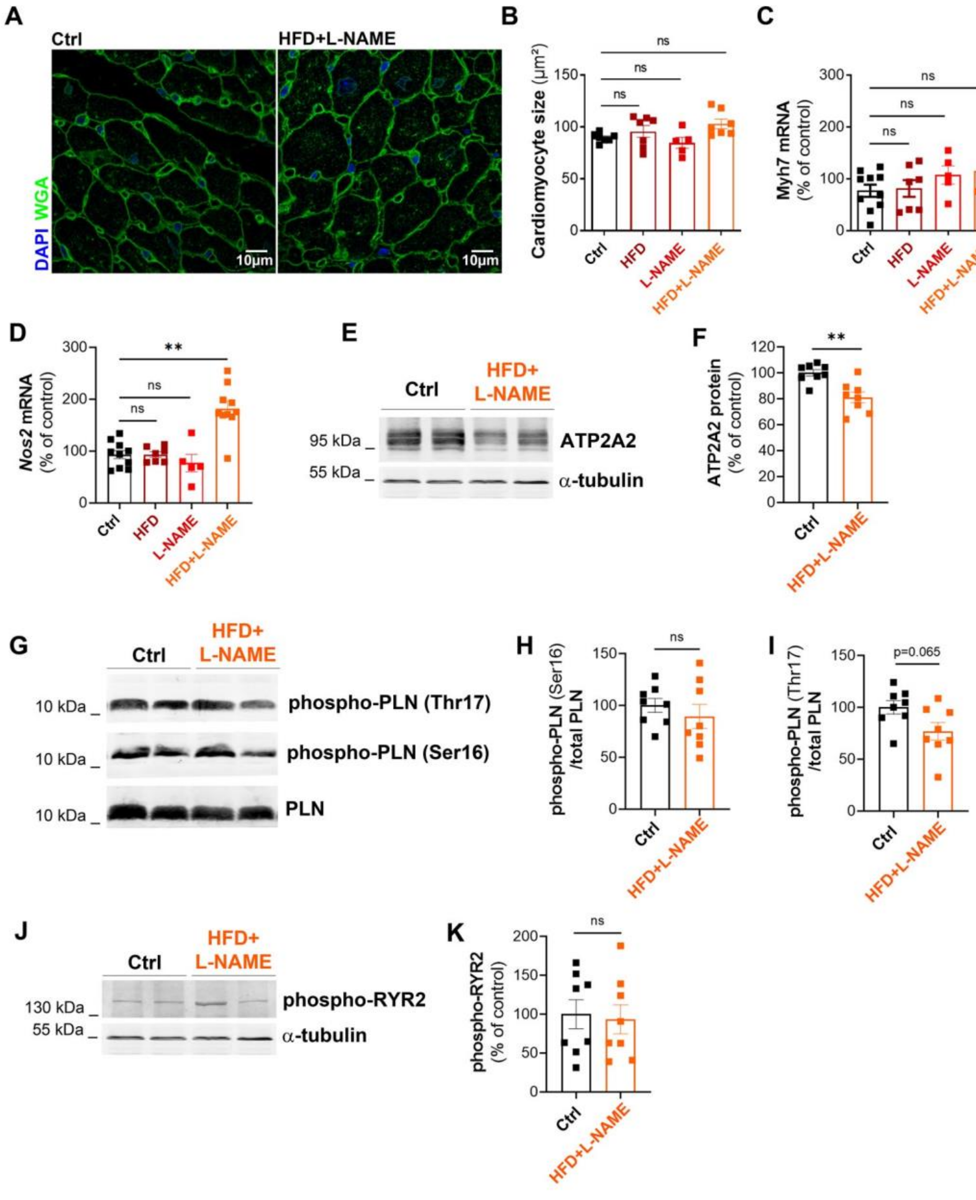

**Supplemental figure 2: OVX female mice submitted to a HFD + L-NAME regimen do not display…**

C57BL/6 female mice were ovariectomized at 7 weeks of age, then submitted or not to a HFD regimen from 8 weeks of age and exposed or not to L-NAME (1 g/L in the drinking water) discontinuously (4 days/week) from 10 week of age. Mice were sacrificed at 20 weeks of age. (**A**) Heart cross sections were stained with FITC-labelled WGA. (**B**) Cardiomyocyte cross section area was measured using Image J software (n=5-7). (**C**) *Myh7* mRNA expression was quantified by RT-qPCR in total heart extract and normalized to Actb mRNA (n=5-10). (**D**) *Nos2* mRNA expression was quantified by RT-qPCR in total heart extract and normalized to Actb mRNA. (**E**) ATP2A2 protein expression was analyzed by western blot in total heart extract and (**F**) quantified using Image J software (n=8). (**G**) PLN phosphorylation was assessed

by western blot analyses and (**H,I**) quantified using Image J software (n=8). (**J**) RYR2 phosphorylation was assessed by western blot analyses and (**K**) quantified using Image J software (n=8).
*: p≤0.05, **: p≤0.01, ***: p≤0.001, ns: not significant (Kruskal-Wallis test with Bonferroni post hoc)

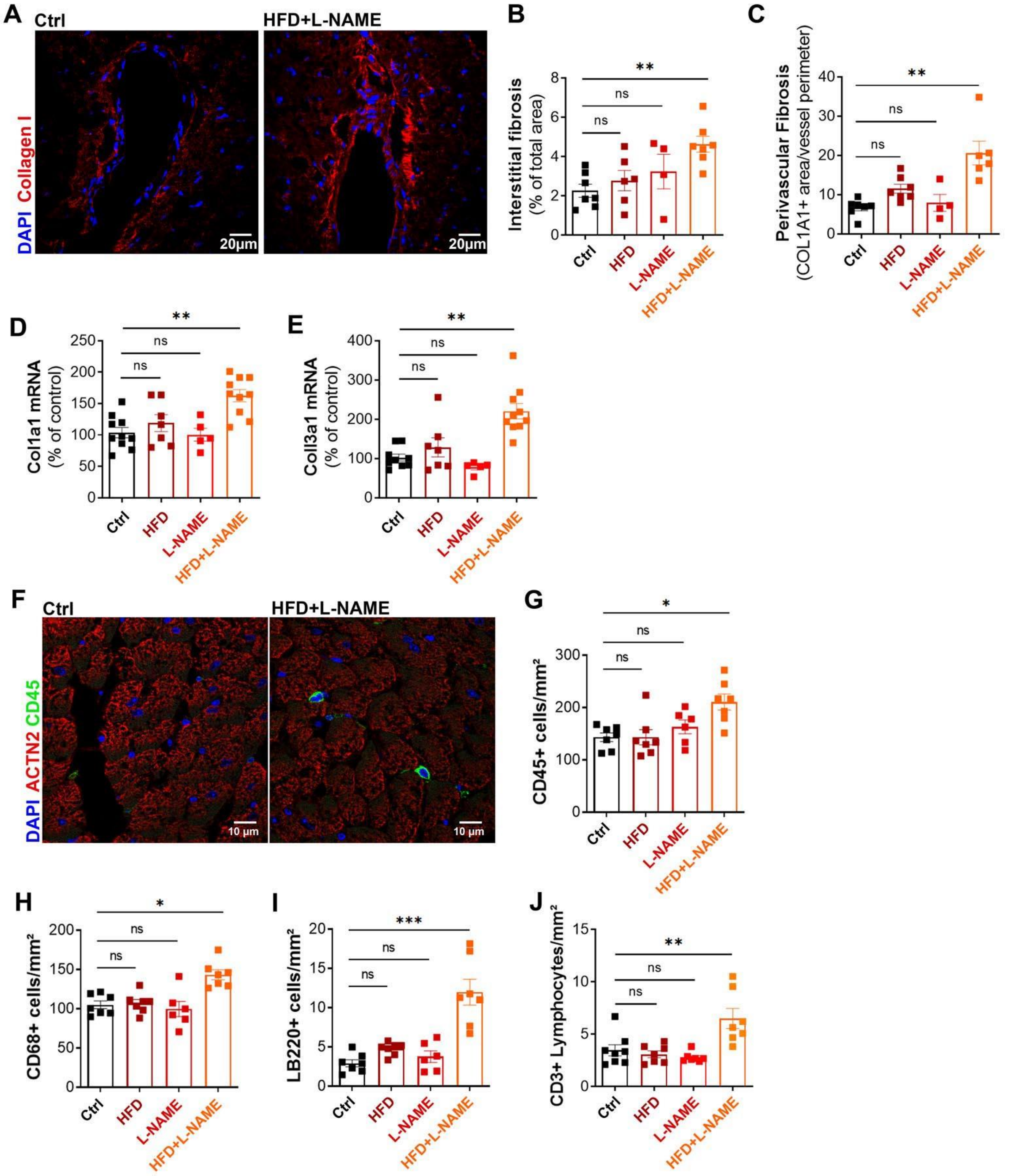

**Supplemental Figure 3: OVX female mice submitted to a HFD + L-NAME regimen display cardiac fibrosis and inflammation.**

C57BL/6 female mice were ovariectomized at 7 weeks of age, then submitted or not to a HFD regimen from 8 weeks of age and exposed or not to L-NAME (1 g/L in the drinking water) discontinuously (4 days/week) from 10 week of age. Mice were sacrificed at 20 weeks of age. (**A**) Heart cross sections were stained with anti-collagen I antibodies to identify fibrosis. (**B**) Interstitial fibrosis was quantified using

Image J software (n=4-7). (**C**) Perivascular fibrosis was quantified using Image J software (n=4-7). (**D**) Col1a1 and (**E**) Col3a1 mRNA expression was quantified by RT-qPCR in total heart extract and normalized to Actb mRNA (n=5-10). (**F**) Heart cross sections were immunostained with anti-CD45 antibodies to identify leukocytes. The number of (**G**) CD45+ leukocytes, (**H**) CD68+ macrophages, (**I**) B220+ lymphocytes and (**J**) CD3+ lymphocytes per $mm^2$ was counted (n=6-7).

*: p≤0.05, **: p≤0.01, ***: p≤0.001, ns: not significant (Kruskal-Wallis test with Bonferroni post hoc)

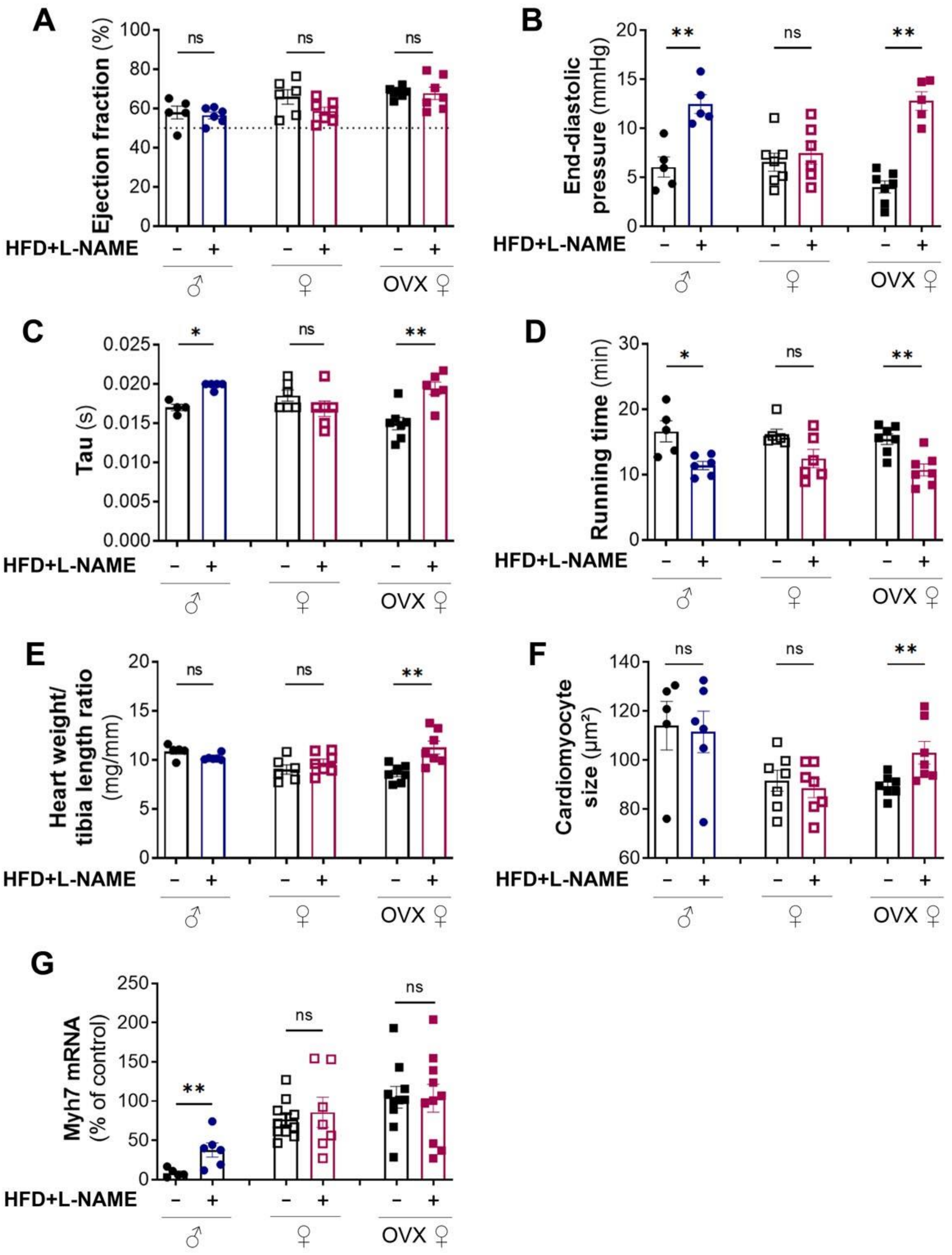

**Supplemental Figure 4: Only male and OVX female mice submitted to a HFD + L-NAME regimen develop diastolic dysfunction.**

C57BL/6 male and female mice were ovariectomized or not at 7 weeks of age, then submitted or not to a HFD regimen from 8 weeks of age and exposed or not to L-NAME (1 g/L in the drinking water) discontinuously (4 days/week) from 10 week of age. Assessments were performed at 20 weeks of age. (**A**) Ejection fraction was assessed via echocardiography (n=5-6). (**B**) LV end diastolic pressure (EDP) was measured by left ventricular catheterization (n=5-6). (**C**) Tau was measured by left ventricular catheterization (n=5-6). (**D**) Running time was assessed on treadmill (n=5-6). (**E**) The heart weight over tibia length was measured (n=5-6). (**F**) Cardiomyocyte cross section area was measured using Image J

software (n=5-6). (G) Myh7 mRNA expression was quantified by RT-qPCR in total heart extract and normalized to Actb mRNA (n=5-10).
*: $p \leq 0.05$, **: $p \leq 0.01$, ***: $p \leq 0.001$, ns: not significant (Mann-Whitney test).

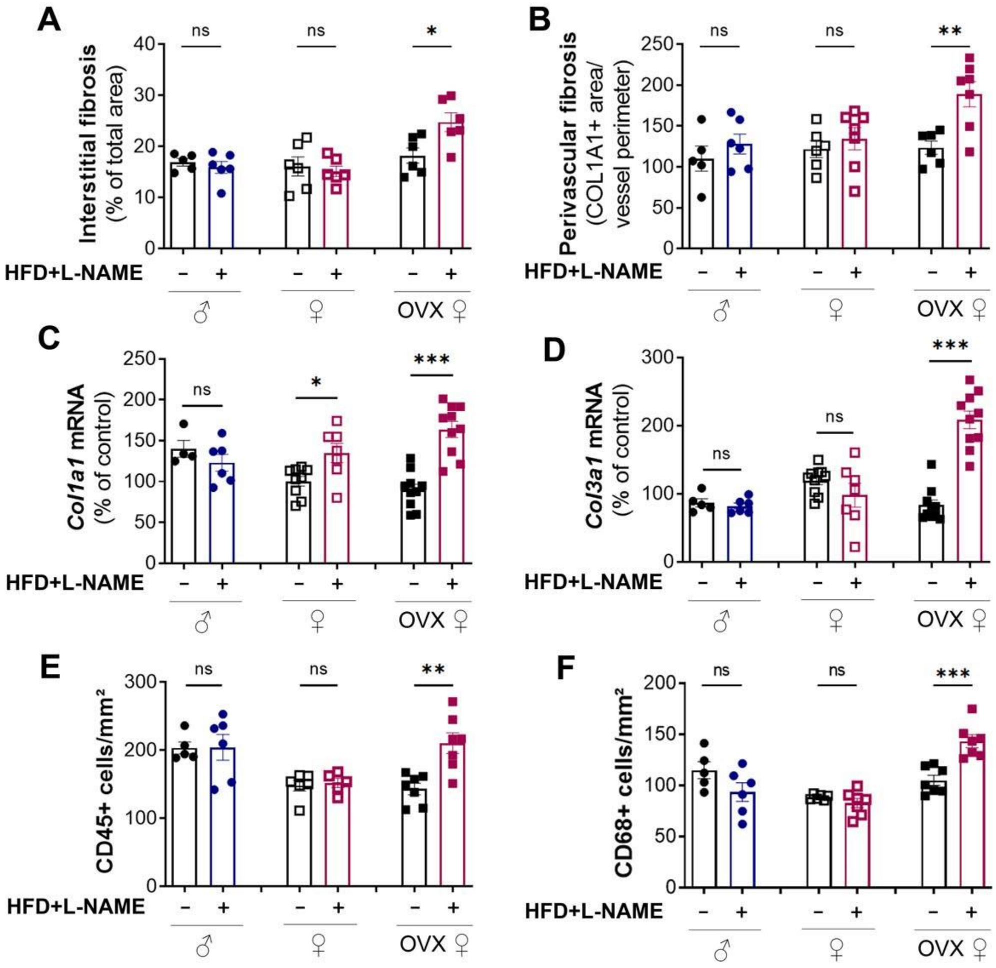

**Supplemental Figure 5: only OVX female mice submitted to a HFD + L-NAME regimen display cardiac fibrosis. Both OVX and non OVX female display cardiac inflammation**

C57BL/6 male and female mice were ovariectomized or not at 7 weeks of age, then submitted or not to a HFD regimen from 8 weeks of age and exposed or not to L-NAME (1 g/L in the drinking water) discontinuously (4 days/week) from 10 week of age. Mice were sacrificed at 20 weeks of age.

(**A**) Interstitial fibrosis was quantified using Image J software (n=5-8). (**B**) Perivascular fibrosis was quantified using Image J software (n=5-8). (**C**) Col1a1 and (**D**) Col3a1 mRNA expression was quantified by RT-qPCR in total heart extract and normalized to Actb mRNA (n=5-10). The number of (**E**) CD45+ leukocytes and (**F**) CD68+ macrophages per mm² was counted (n=5-10).

*: p≤0.05, **: p≤0.01, ***: p≤0.001, ns: not significant (Mann-Whitney test)

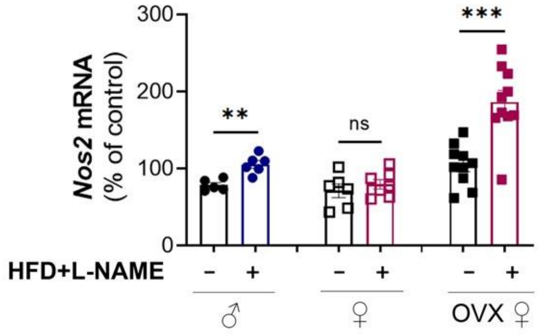

**Supplemental Figure 6: Diastolic dysfunction is associated with increased Nos2 levels in both males and females**

C57BL/6 male and female mice were ovariectomized or not at 7 weeks of age, then submitted or not to a HFD regimen from 8 weeks of age and exposed or not to L-NAME (1 g/L in the drinking water) discontinuously (4 days/week) from 10 week of age. Nos2 mRNA expression was quantified at in 20 week old mice by RT-qPCR in total heart extract and normalized to Actb mRNA (n=5-10).
